# Retrovirus-derived acquired genes, *RTL5* and *RTL6*, are novel constituents of the innate immune system in the eutherian brain

**DOI:** 10.1101/2021.12.29.474483

**Authors:** Masahito Irie, Johbu Itoh, Ayumi Matuszawa, Masahito Ikawa, Toru Suzuki, Yuichi Hiraoka, Fumitoshi Ishino, Tomoko Kaneko-Ishino

## Abstract

Retrotransposon Gag-like 5 (*RTL5*, also known as sushi-ichi-related retrotransposon homolog 8 (*SIRH8*)) and *RTL6* (aka *SIRH3*) are eutherian-specific genes presumably derived from a retrovirus and phylogenetically related to each other. *RTL5* encodes a strongly acidic protein while *RTL6* encodes an extremely basic protein, and the former is well conserved and the latter extremely well conserved among the eutherians, indicating their unique and critically important roles as acquired genes. Here we report that *RTL5* and *RTL6* are microglial genes playing roles in the front line of brain innate immune responses against distinct pathogens. Venus and mCherry knock-in mice exhibited expression of RTL5-mCherry and RTL6-Venus fusion proteins in microglia and as extracellular granules in the central nervus system (CNS), and displayed a rapid response to pathogens such as lipopolysaccharide (LPS), double-stranded (ds) RNA analog and non-methylated CpG DNA. These proteins trapped pathogens in microglia in a variety of RTL-pathogen complexes depending on the pathogens. These results demonstrate that *RTL5* and *RTL6* exert functional effects against different hazardous substances cooperatively and/or independently to protect the developing and/or mature brain. This provides the first evidence that retrovirus-derived genes play a role in the innate immune system of the eutherian brain.

## Introduction

In humans and mice, 11 retrotransposon Gag-like (*RTL*) genes encode proteins exhibiting 20∼30 % homology to the sushi-ichi long terminal repeat (LTR) retrotransposon GAG proteins, and in some cases also to POL proteins. They exhibit a variety of biological functions in the eutherian developmental system and each protein has unique amino acid (aa) sequence and length from 112 to 1,744 aa residues (Brandt et al., 2005; Yongson et al., 2005; Ono et al., 2006; Kaneko-Ishino and Ishino, 2012, 2105). These are good examples of exaptation, that is, gaining novel function(s) for specific purposes during the course of evolution, as originally proposed by Gould and his colleagues (Gould and Vrba, 1982; Brosius and Gould. 1992): *Peg10* (*aka Rtl2* or *Sirh1*), *Rtl1* (*aka Peg11* or *Sirh2*) and *<u>L</u>eucine zipper, <u>d</u>ownregulated in <u>c</u>ancer 1* (*Ldoc1, aka Rtl7* or *Sirh7*) play essential but distinct roles in the formation, maintenance and endocrine regulation of the placenta in mice, respectively (Ono et al., 2006; Sekita et al., 2008; Kagami and Sekita et al., 2008; Kaneko-Ishino and Ishino, 2012; Naruse et al., 2014; Kaneko-Ishino and Ishino, 2105). *Rtl1* also is involved in fetal/neonatal muscle development (Kitazawa et al., 2020) as well as in the functions of the corticospinal tract and corpus callosum, mammalian- and eutherian-specific brain structures (Kitazawa et al., 2021), respectively. In addition, *Rtl4* (*aka Sirh11*) is related to cognitive function in the brain via noradrenaline regulation (Irie et al., 2015). *RTL6* (aka *SIRH3* or *LDOC1-like* (*LDOC1L*)) is of great interest because it is extremely conserved in eutherians. In this work, we address how *RTL6* and the phylogenetically related *RTL5* (also called *SIRH8* or retrotransposon Gag domain like 4 (*RGAG4*)) contribute to the present day eutherian innate immune system as eutherian-specific, acquired genes.

Microglia originate from extraembryonic yolk sac in early development, migrate to the embryo and settle in the brain in the fetal stage, then ultimately propagate throughout the brain over the course of life (Ginhoux et al., 2010, 2013). Microglia are the primary innate immune cells of the brain and play a central role in the immune responses to various pathogens via a variety of Toll-like receptors (TLRs) (Hanisch and Kettenmann, 2007; Norris and Kipnis, 2018). Moreover, in the neonatal brain microglia are involved in shaping neuronal circuits during development via regulating neurogenesis. They induce filopodia formation by direct contact with neurons and phagocytose supernumerary or unwanted synapses as well as pruning excess astrocytes in the developing amygdala (Hanisch and Kettenmann, 2007; Sierra et al., 2010; Reemst et al., 2016).

## Results

### *RTL5 and RTL6* are eutherian-specific genes encoding a strongly acidic and extremely basic protein, respectively

Based on the genome data, *RTL5* is located at the end of intron 1 of Nance-Horan syndrome like 2 (*NHSL2*) in the opposite direction (Figs. 1a and b), while *RTL6* lies between *Shisa like 1* (*SHISAL1*) and Proline rich 5 (*PRR5*) (Fig. 1c). These locations are conserved in all four eutherian lineages, but no orthologues exist in birds, monotremes or marsupials.

**Figure 1.**
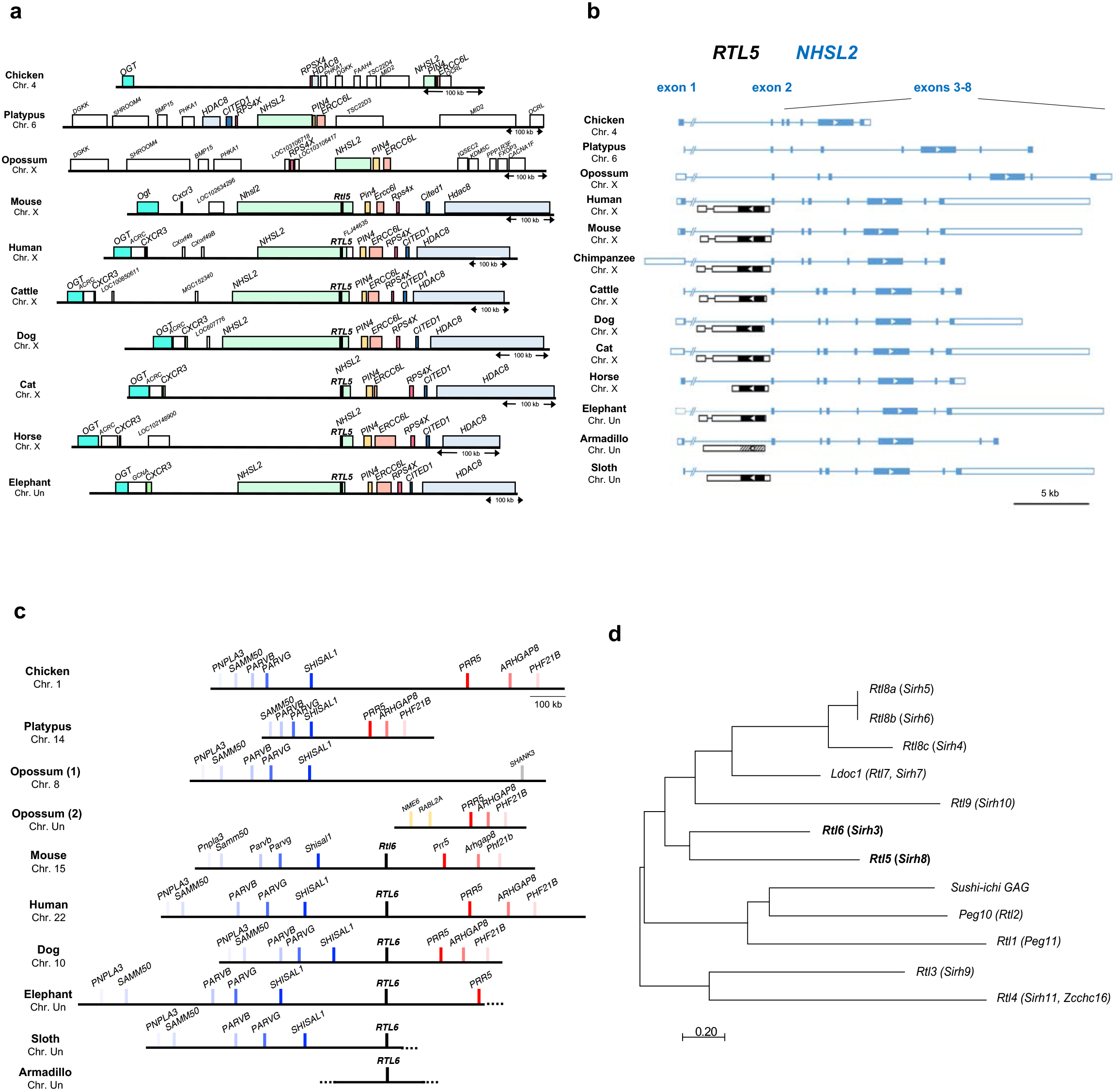
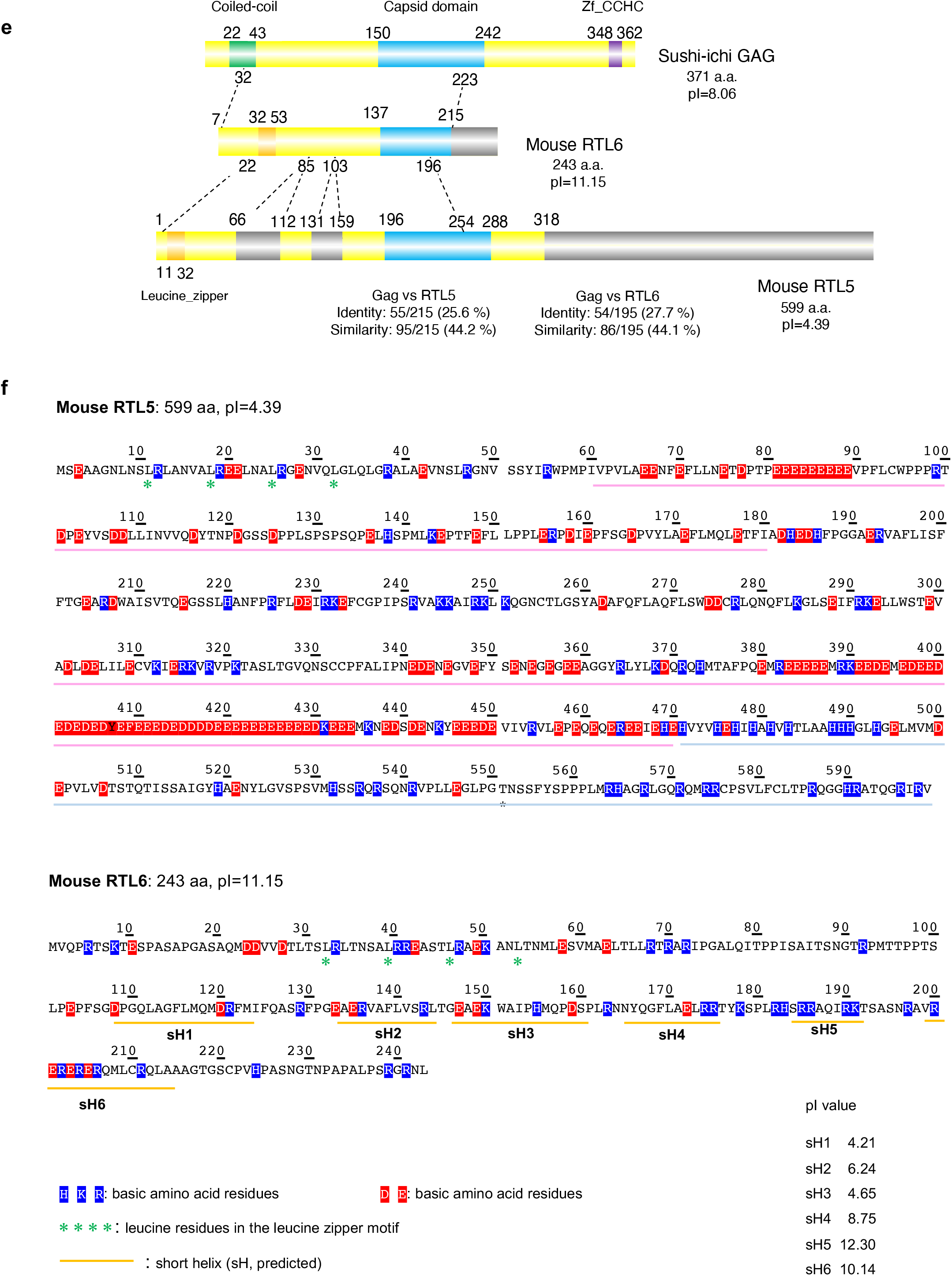
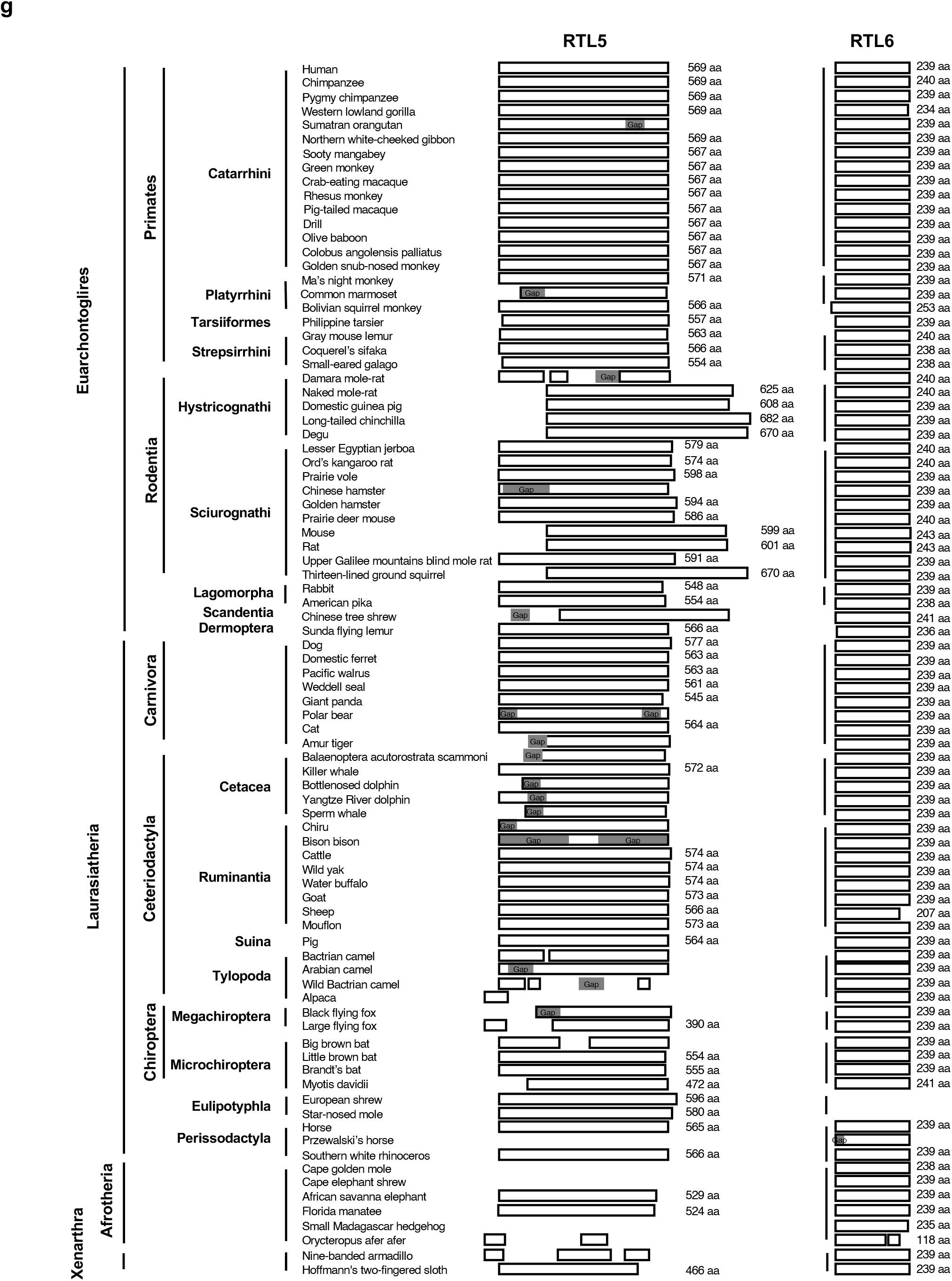
Conservation of the *RTL5* and *RTL6* genes in eutherians and characters of RTL5 and RTL6 proteins. **(a)** Chromosomal location of *RTL5* across the eutherians. The chromosomal region around *NHSL2* where *RTL5* is located is conserved in eutherians but slightly different in marsupials, monotremes and birds except for *NHSL2-PIN4-ERCC6*, suggesting that chromosome rearrangements occurred just around the *RTL5* insertion site in a lineage-specific manner. The boxes in identical colors represent orthologous genes. **(b)** The chromosomal locations of *RTL5* in the intron 1 of *NHSL2*. **(c)** The chromosomal locations of *RTL6* between *SHISAL1*and *PRR5*. The boxes in identical colors represent orthologous genes. In marsupials, *SHISAL1* and *PPR5* are separated into two different chromosomal loci, suggesting that chromosomal rearrangement occurred just around the *RTL6* insertion site. **(d)** A phylogenic tree of sushi-ichi *GAG* and all the mouse *RTL* genes. The scale bar indicates the number of substitutions per site. **(e)** Alignment of the sushi-ichi GAG (top), mouse RTL6 (middle) and RTL5 (bottom) proteins. Coiled-coil and leucine zipper motifs in the N-terminus, a capsid domain in the middle and a zinc finger CCHC domain in the C-terminus are represented as green, orange, blue and purple boxes, respectively. The grey boxes indicate they exhibit no homology with the GAG protein. **(f)** Amino acid sequences of the mouse RTL5 and RTL6 proteins, along with the number and position of their basic and acidic amino acid residues. The mouse RTL5 protein contains 72 basic and 137 acidic amino acid (aa) residues out of a total of 599 aa, and so is strongly acidic (pI=4.39), especially the two regions underlined in pink, from 61 to 180 aa (a total of 120 aa: pI=3.55) and from 301 to 470 aa (a total of 170 aa: pI=3.82). However, they have also a basic part underlined in light blue, from 471 to 599 aa (a total of 129 aa: pI=10.02) in the C-terminus. The mouse RTL6 protein contains 34 basic and 20 acidic aa residues out of a total of 243 aa, and thus is extremely basic (pI=11.15). In particular, the basic aa are concentrated in the short C-terminus helix (sH) 5 and sH6 (pI=12.30 and 10.14, respectively) **(g)** Conservation of the RTL5 and RTL6 proteins in the eutherians. Among the 86 eutherian species, *RTL5* has been confirmed in 82. It is likely that *RTL5* has become dysfunctional in certain eutherian species. There is no data published on the Prairie vole, Przewaiski’s horse, Cape golden mole or Cape elephant shrew. Among the 86 eutherian species, *RTL6* was confirmed in 84 species. The RTL6 protein is mostly comprised of 239 to 243 amino acids with two notable exceptions, sheep (207 aa) and Orycteropus afer afer (168 aa). There is no data available on the European shrew or Star-nosed mole.

Phylogenetically, *RTL5* and *RTL6* are the most closely related among the *RTL*s (Fig. 1d): the RTL5 and RTL6 proteins have a leucine-zipper motif in their N-terminus and exhibit a high degree of similarity to each other (Fig. 1e) as well as to the suchi-ichi GAG protein (44.2 and 44.1% similarity and 25.6 and 27.7% identity, respectively). The mouse RTL5 protein comprising 599 aa is strongly acidic (pI=4.39), while the mouse RTL6 protein comprising 243 aa is extremely basic (pI=11.15) (Fig. 1f). *RTL5* is evolutionary well conserved in eutherians (dN/dS ratio = 0.3∼0.5) (Table 1, top), but may be functionally inactive in some species due to a variety of mutations (Fig. 1g, left). In contrast, *RTL6* exhibits an extremely low dN/dS ratio (mostly < 0.05) across eutherian species (Table 1, bottom), indicating that *RTL6* has been subjected to quite strong purifying selection during the course of evolution (Fig. 1g, right).

**Table 1.**
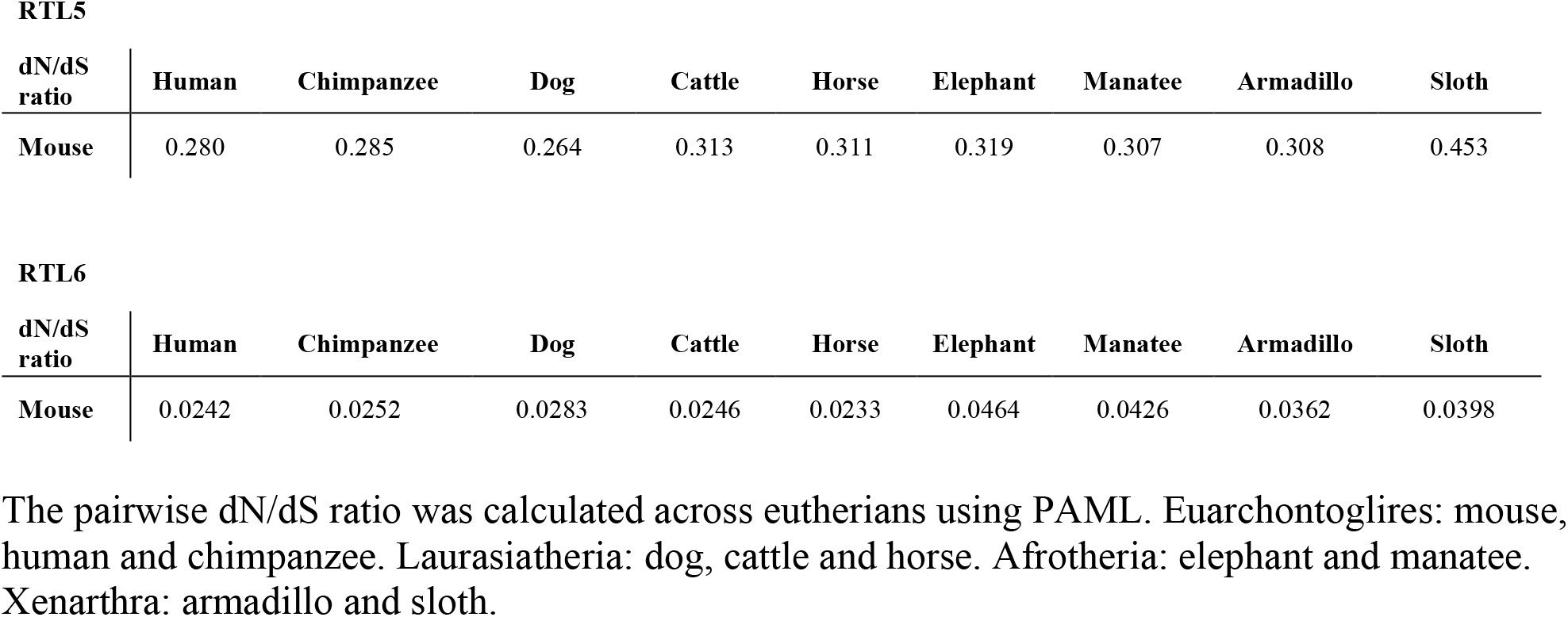
dN/dS ratio of *RTL5* and *RTL6* between mouse and nine other eutherian species.

### RTL6 protein expression *in vivo*

First, we focused on the characterization of *Rtl6* in mice, because despite the extremely conserved nature of the RTL6 protein, mouse RTL6 is encoded in the fourth ORF in a reference sequence (RefSeq) of the *Rtl6* transcript in GenBank (Fig. 2a, top panel, the second line). This strongly suggests that expression of the mouse RTL6 protein is quite low even if clearly expressed, so there may be a large discrepancy between the expression levels of the *Rtl6* mRNA and RTL6 protein (see next section). Actually, it was very difficult to detect the mouse RTL6 protein in any tissues and organs by usual methods, such as Western blotting or immunostaining analysis using either commercially available anti-RTL6 antibodies or those of our own making.

**Figure 2.**
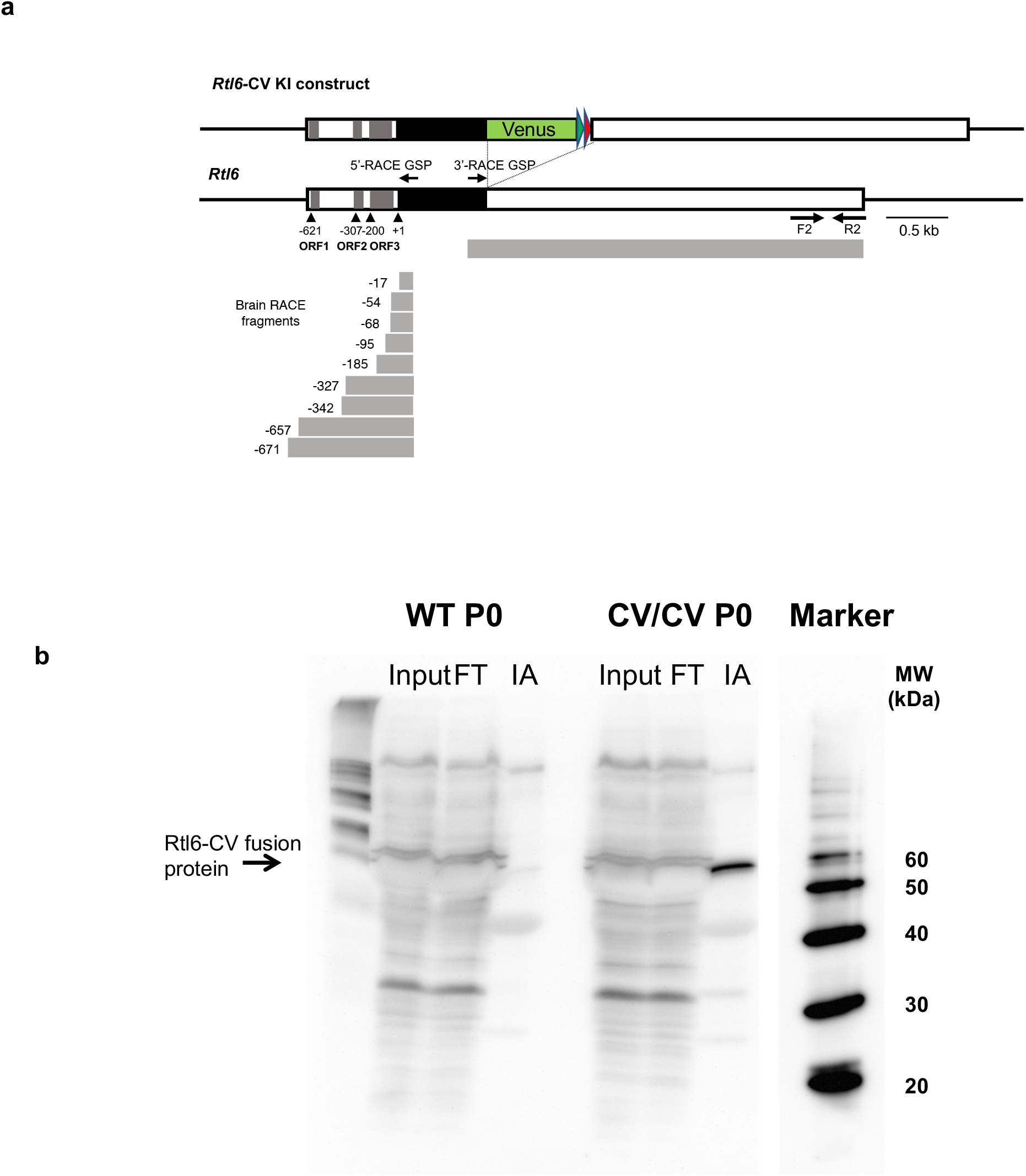
Confirmation of RTL6 protein expression by RTL6-Venus fusion protein. **(a)** The conserved mouse RTL6 ORF is encoded in the fourth ATG in the Refseq (the second line). The 5’- and 3’-RACE fragments (grey boxes) are presented below. The four black triangles represent ATG codon sites. The construction of the *Rtl6-Venus* KI mouse is also presented (the first line). Red and blue triangles represent loxP and Frt sites, respectively. **(b)** Expression of the RTL6-Venus fusion protein in neonatal brain of *Rtl6*-CV KI mice. Immunoaffinity experiment. Input: total extract of P0 brain. FT: the flow through fraction, IA: the elution fraction from anti-Venus antibody beads. MW: Molecular weight marker. The RTL6-CV proteins are indicated with an arrow. The RTL6-Venus protein was only detected in the *Rtl6*-CV samples (right) but not in the wild type control (left).

A 5’-RACE experiment using the 8 week (w) adult brain with gene specific primers (GSPs) from inside of the protein coding region demonstrated that among nine different 5’-RACE fragments, five of them, starting at the -185, -95, -68, -54 and -17 nucleotide positions from ATG (+1), respectively, have RTL6 as the first ORF, suggesting that the RTL6 ORF can be expressed from such short mRNAs (Fig. 2a, bottom). Then we generated Venus KI mice in which a Venus ORF is integrated into the endogenous *Rtl6* locus just after its C-terminus (*Rtl6*-CV strain) to detect the mouse RTL6 protein *in vivo* in the same tissues and organs as the endogenous RTL6 protein (Fig. 2a, top panel, the first line). The RTL6-Venus fusion protein was detected by immunoaffinity (IA) chromatography using an anti-Venus (GFP) antibody with the expected molecular weight of 54 kDa (27 kDa each for the RTL6 and Venus proteins) in the neonatal day 0 (P0) KI mouse brain (Fig. 2b), demonstrating that it was evidently expressed in the brain.

### RTL6 expression in microglia in the CNS

The RTL6-Venus protein is exclusively expressed in microglia and also is present as extracellular granules in the brain. To obtain the precise positional as well as the relative fluorescent strength information using confocal fluorescence microscopy, it is essential that the target Venus signal (emission peak at 530 nm) be separated from various kinds of autofluorescence (Af) in the embryo, brain and other tissues (Fig. 3a). Therefore, the data processing function known as Multi-channel Unmixing or Automatic Composition Extraction (ACE) was applied. Confocal fluorescence microscopy analysis of the *Rtl6*-CV mice demonstrated that its expression was observed to be weak in the yolk sac and embryo on embryonic day 9.0 (E9.0) (weaker than the 8^th^ strongest signal (ACE8)), then became very strong (ACE2) in the CNS by E13.5, and its highest expression was observed in the period of the perinatal and postnatal day 0∼3 brain (P0∼3, ACE1∼2) (Fig. 3a). It is thus concluded that the RTL6 protein is exclusively expressed in the CNS in mice (Fig. 3b), even though the qPCR experiment demonstrated that *Rtl6* mRNA expression was higher in the muscle, kidney and testis than the brain in 4 and 8w adults (Fig. 3c).

**Figure 3.**
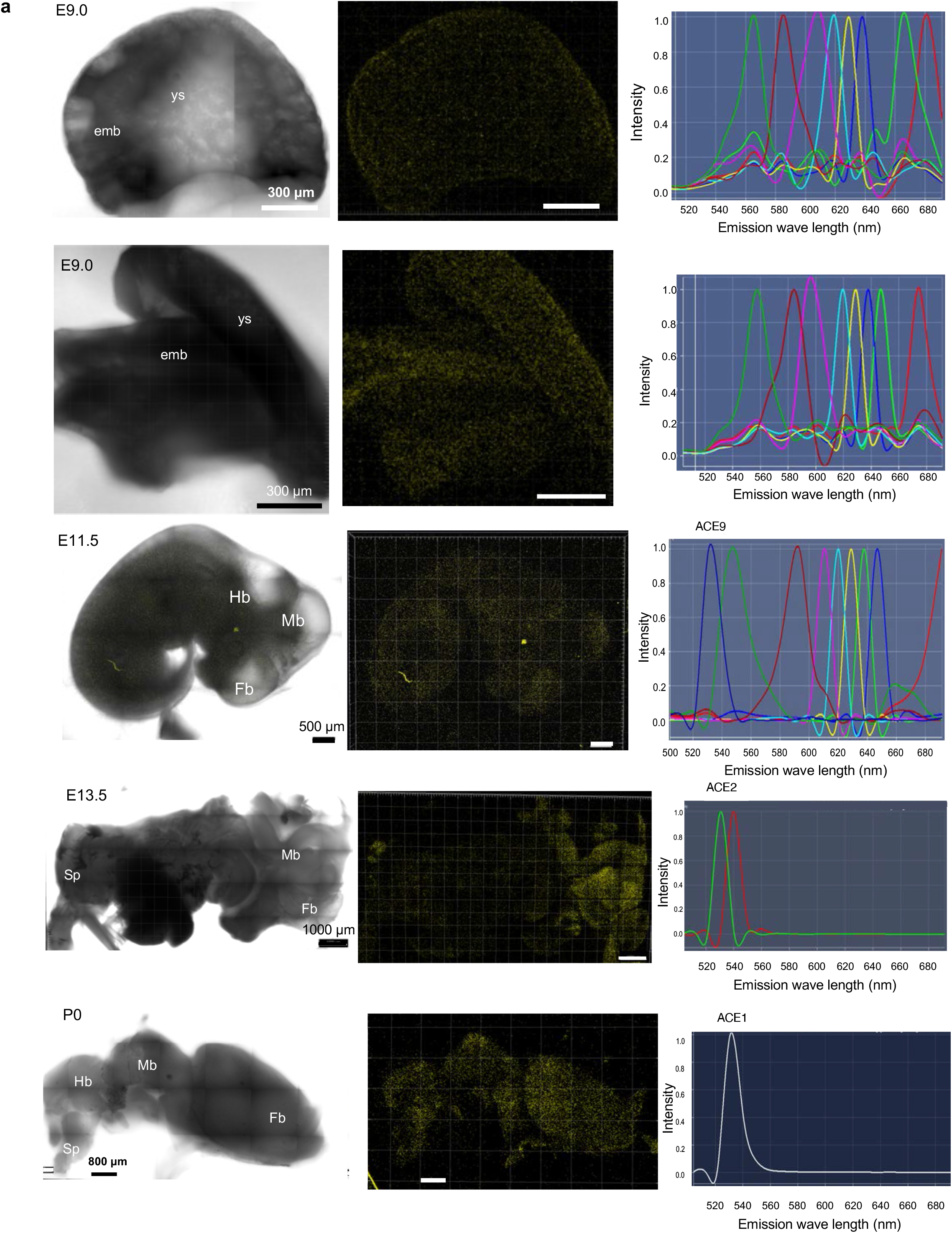
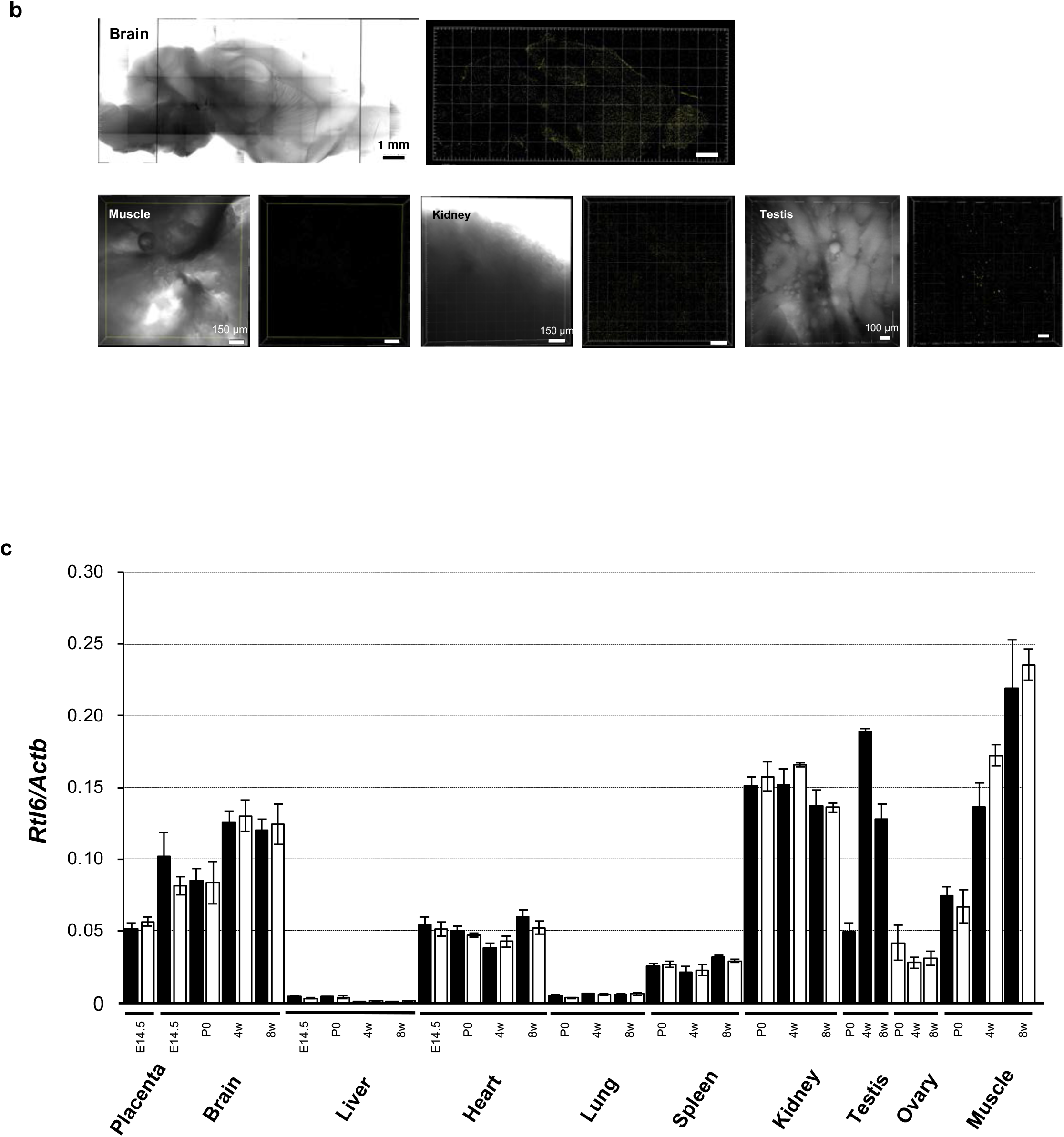
RTL6 protein expression during mouse development. **(a)** RTL6-Venus signals detected from the early developmental stage to the neonatal stages. Left: transmission image. Middle: Venus signals (530 nm). Right: ACE spectrum profiles. Top and second: E9.0 embryo and yolk sac. Venus signal was detected in both embryos and yolk but very weak (out of ACE8, N>5). Third and fourth: Venus signal was still very weak (ACE9) in E11.5 embryo (N>3) but became strong in the CNS in E13.5 embryo (ACE2) (N>3). Bottom: P0 brain exhibited the strongest Venus signal (ACE1) (N>3). **(b)** The Venus signals in the brain (top) in 6w adults (N=4) and hindlimb muscle (bottom left), kidney (bottom middle) and and testis (bottom right) in 4w adults (N=2). Transmission (left) and Venus signal images (right) are shown in each column. In these organs and tissues, no Venus signals were detected in the top range of ACE9 signals. However, in the testis, a few Venus-positive large granules (5 μm in diameter) were observed in the seminiferous tubules. **(c)** The expression levels of *Rtl6* mRNA. *Rtl6* mRNA expression level in several organs and tissues during development. *Rtl6* mRNA expression in the muscle, kidney and testis is higher than that in the brain during development and growth. Black and white bars: male and female samples, respectively.

Venus-positive cells were observed in the hypothalamus and olfactory bulb regions on the internal surface of the brain in neonates (Fig. 4a). They were completely merged with cells expressing a microglial marker, anti-ionized calcium-binding adapter molecule 1(Iba1) (Utan et al. 1995; Ito et al., 1998) (Fig. 4b), demonstrating that the RTL6-Venus protein is expressed exclusively in microglia with a variety of morphologies, such as round, ameboid, irregular and elongated shapes together with processes (Hanisch and Kettenmann, 2007; Norris and Kipnis, 2018). In contrast, in internal regions such as the hippocampus (Fig. 4c, top and second) and amygdala (Fig. 4d, top and middle), most of the Venus signals were spread out as small extracellular granules (much less than 1 μm in diameter), while Venus-positive cells were only rarely observed. In the hippocampus, the small Venus-positive dots were accumulated alongside the pyramidal neurons within the hippocampal sub-regions 1∼3 (CA1∼CA3) and granular cells in the dentate gyrus (DG) (Fig. 4c, top and second), where only a small number of Venus-positive cells were detected (Fig. 4c, third and bottom). In addition, larger extracellular granules (1∼3 μm in diameter), each comprised of small extracellular dots, frequently observed in the amygdala and certain neonatal midbrain regions, such as the inferior colliculus and substantia nigra (Fig. 4d, bottom).

**Figure 4.**
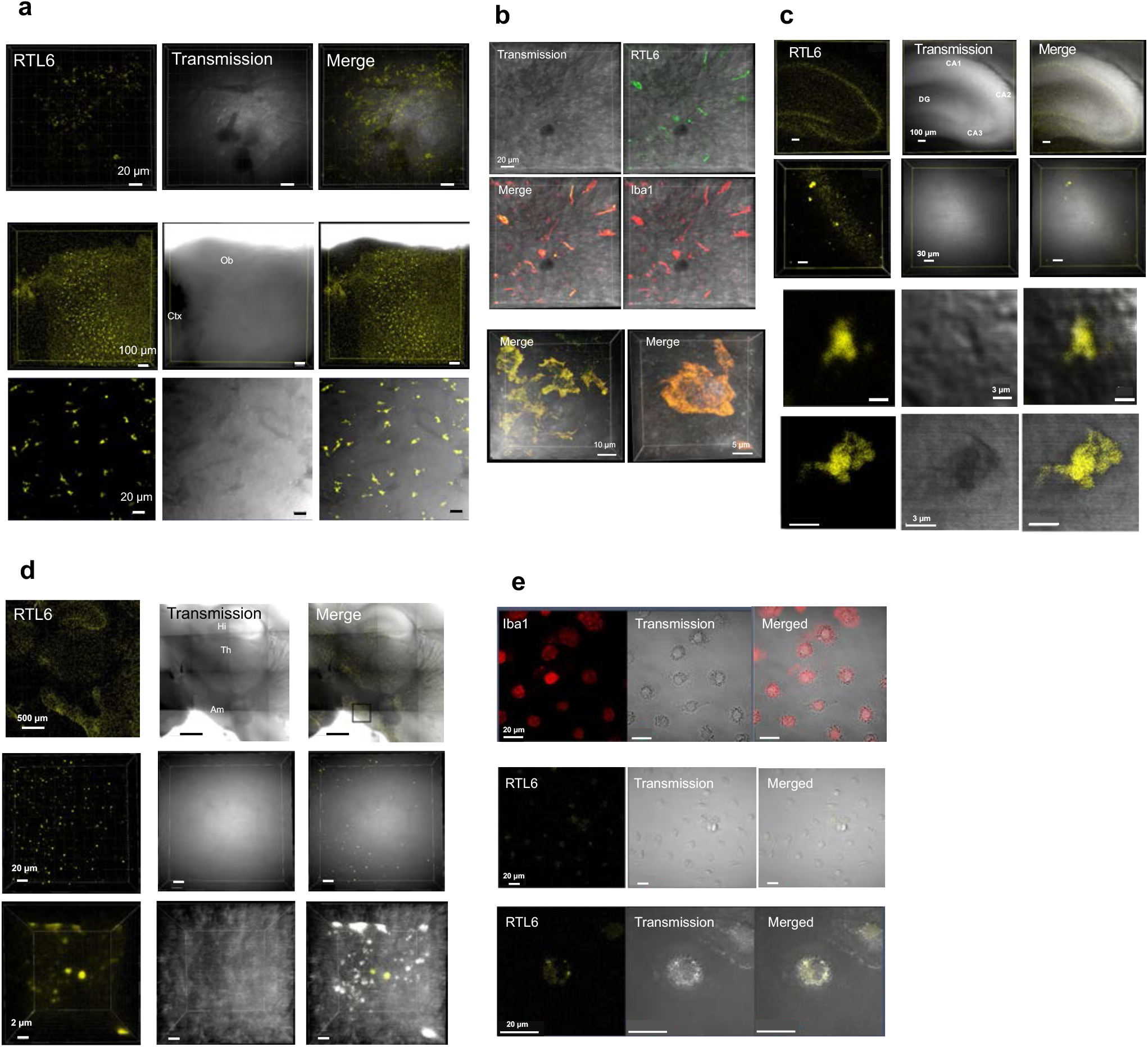
RTL6 expression in microglia. **(a)** Venus-positive cells in the hypothalamus (top), olfactory bulb (middle) and enlarged images of the olfactory bulb region (bottom). Ctx: cerebral cortex. Ob: olfactory bulb. **(b)** Iba1staining of the Venus-positive cells in the hypothalamus (top and middle, see also Supplementary movie 1) and enlarged images of the Venus-positive cells (bottom, see also Supplementary movies 2 and 3). **(c)** Adult hippocampus (6w). Top: the dorsal part of the hippocampus. Second: enlarged pictures of the CA2 region. Third and bottom: Venus positive cells in the DG region. CA1∼3: cornu ammonis subregions 1-3. DG: dentate gyrus. **(d)** Top: the posterior half of the 6w brain. Middle: adult amygdala (6w) (enlarged pictures of the black square in the top right figure). Bottom: enlarged pictures of Venus-positive large granules in the amygdala region. The autofluorescence signals are merged with the Venus signals but their strength is different among the granules. Am: amygdala, Hi: hippocampus. Th: thalamus. **(e)** Isolated microglial cells from P1 neonatal brain. Top: Iba1 staining. Middle: RTL6 expression. Bottom: enlarged picture showing RTL6 granules in microglia.

Finally, we isolated the microglial cells from the P1 neonatal KI brain and cultured them *in vitro* (Folden and Combs, 2007; Lian et al. 2016). The cells that were collected from the cultured Petri dishes by tapping had a round shape or an amoeba-like morphology (Fig. 4e, top) and at least 40 % of the P1 microglial cells actually expressed the RTL6-CV protein at a relatively high level (Fig. 4e, middle) as intracellular granules, like the Venus-positive cells in the brain (Fig. 4e, bottom).

### RTL5 is also expressed in microglia

RTL5 is also expressed in microglia like RTL6. Using the *Rtl5*-mCherry KI (*Rtl5*-CmC) mice (Fig. 5a) and the *Rtl6*-Venus and *Rtl5*-mCherry double KI (DKI) mice generating by mating these strains, we confirmed RTL5 expression in the microglia in the hypothalamus and olfactory bulb on the inner surface of the hemispheres (Figs. 5b, c and d). We confirmed RTL5 expression in isolated and cultured microglial cells (Fig. 5b). An autofluorescence that peaked at 610 nm was frequently detected in many tissues and organs (also in microglia), but the mCherry (610 nm) signal was usually distinguishable by using the ACE system (Figs. 5b and e, Table 2). The RTL5-positive cells were detected in the hypothalamus (Fig. 5d, top) and RTL5 coexisted with RTL6 in the same microglial cells at different ratios, but its intensity was usually lower than that of RTL6 (Fig. 5d, bottom). The RTL5 protein was present as granules in the microglia in the olfactory bulb (Fig. 5f, top) and also detected in the microglia in the cerebral cortex with a round morphology a few processes (Fig. 5f, bottom).

**Table 2.** Separation of mCherry signals from autofluorescent (610 nm) signals in the isolated microglia.

|  | WT |  |  |  | RTL5-CC |  |  |  |  |  |  |
| --- | --- | --- | --- | --- | --- | --- | --- | --- | --- | --- | --- |
|  | 1 | 2 | 3 | 4 | 1 | 2 | 3 | 4 | 5 | 6 | 7 |
| mCherry | 39 | 82 | 21 | 61 | 721 | 749 | 608 | 802 | 2376 | 3599 | 3829 |
| af610 | 1740 | 3383 | 1358 | 1995 | 142 | 157 | 157 | 163 | 477 | 712 | 633 |
| mCherry/af610 | 0.022 | 0.024 | 0.015 | 0.031 | 5.08 | 4.77 | 3.87 | 4.92 | 4.98 | 5.05 | 6.05 |

Autofluorescence-derived 610 nm and mCherry signals were counted in single microglial cells from WT (N=4) and *Rtl5-mCherry* KI mice (N=7). The mCherry signals detected in the WT microglia seemed experimental error, but negligible compared with those from microglia in the *Rtl5-mCherry* KI mice, indicating that these two signals could be separated each other in this experimental system.

**Figure 5.**
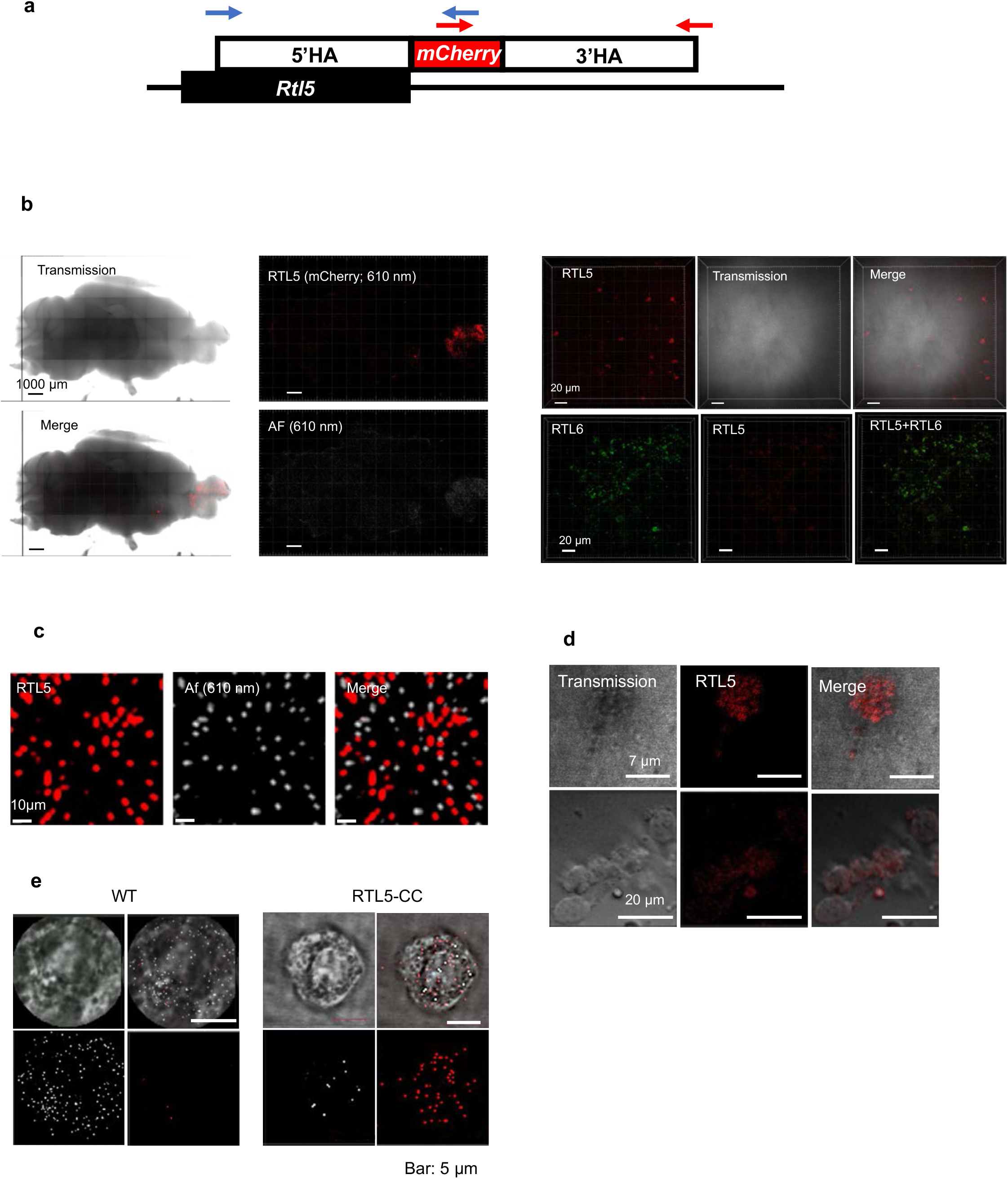
RTL5 expression in microglia. **(a)** The construction of the *Rtl5-mCherry* KI mouse. **(b)** Isolated microglial cells from P1 neonatal brain of wild type (left) and *Rtl5-mCherry* KI mice (right). Top left: transmission image. Top right: merged image. Bottom left: Af610nm-derived signals. Bottom right: mCherry-derived signals (see also Table 2). **(c)** RTL5 expression on the inner surface of the hemispheres in 4w adults (N=2). The pictures of mCherry (top, 610 nm, red signals) and Af610nm (bottom, white signals) are presented. **(d)** mCherry-positive cells in the hypothalamus (top). The mCherry-signals were lower than the Venus-signals in the same microglial cells (bottom). **(e)** Enlarged views of the olfactory bulb. Most of the mCherry-derived red signals were not merged with the Af610nm-derived white signals, indicating they recognized different substances. **(f)** mCherry-positive cells in the olfactory bulb (top) and cerebral cortex (bottom).

### Function of RTL5 and RTL6 against certain pathogen

RTL6 and RTL5 quickly reacted to certain pathogens, including lipopolysaccharide (LPS), dsRNA and non-methylated dsDNA. They formed different RTL-pathogen complexes in microglia depending on the pathogens used. Microglia are the primary innate immune cells of the brain and play a central role in the immune responses to various pathogens via a variety of Toll-like receptors (TLRs) (Fiebich et al., 2018). Therefore, we analyzed the response of RTL6 and RTL5 proteins to pathogens in the brain.

Five to 10 min after the injection of Alexa594-labeled-LPS (emission peak 617 nm), fresh brain was dissected out and directly examined under confocal fluorescence microscopy for 1∼2 hrs or after fixation with paraformaldehyde (PFA). On lower magnification, the RTL6 (green) signal was observed to be accumulated in the LPS injected regions (shown in artificial blue) (Fig. 6a). Despite a methodological limitation in obtaining the absolute value of each signal intensity, we were able to calculate the relative intensity of each signal in the whole brain from the left (olfactory bulb) to right (cerebellum) sides (Fig. 6a, a red line in the left middle panel), indicating that mainly RTL6 and to a lesser extent RTL5 accumulated at the LPS injection sites in proportion to the LPS amount (Fig. 6b). On higher magnification, the microglia that had accumulated near the blood capillaries in the cerebral cortex had transformed into giant flattened cells and formed a barrier-like structure along with the blood capillaries by assembling together (Fig. 6c, top panels). It is known that bacterial infections and/or inflammation induce similar multinuclear giant cells (MNGCs) comprised of microglia (Peterson et al. 1996; Hornik et al., 2014), but these flattened cells were not fused because a clear cellular boundary was evident (Figs. 6c, transmission images in top and bottom panels, and 6d, top and middle panels). LPS was incorporated along the cellular edges where RTL6 had accumulated on the cytoplasmic side in the flattened microglia (Fig. 6c, top panels). In addition, a large RTL5/RTL6/LPS complex frequently appeared at the intersection between three or four giant flattened cells (Figs. 6c, bottom panels, and 6d, top panels). A sequence of photographs at 0.7 μm interval suggests that RTL6 formed the complex core and trapped LPS on its surface (Fig. 6d, bottom panels).

**Figure 6.**
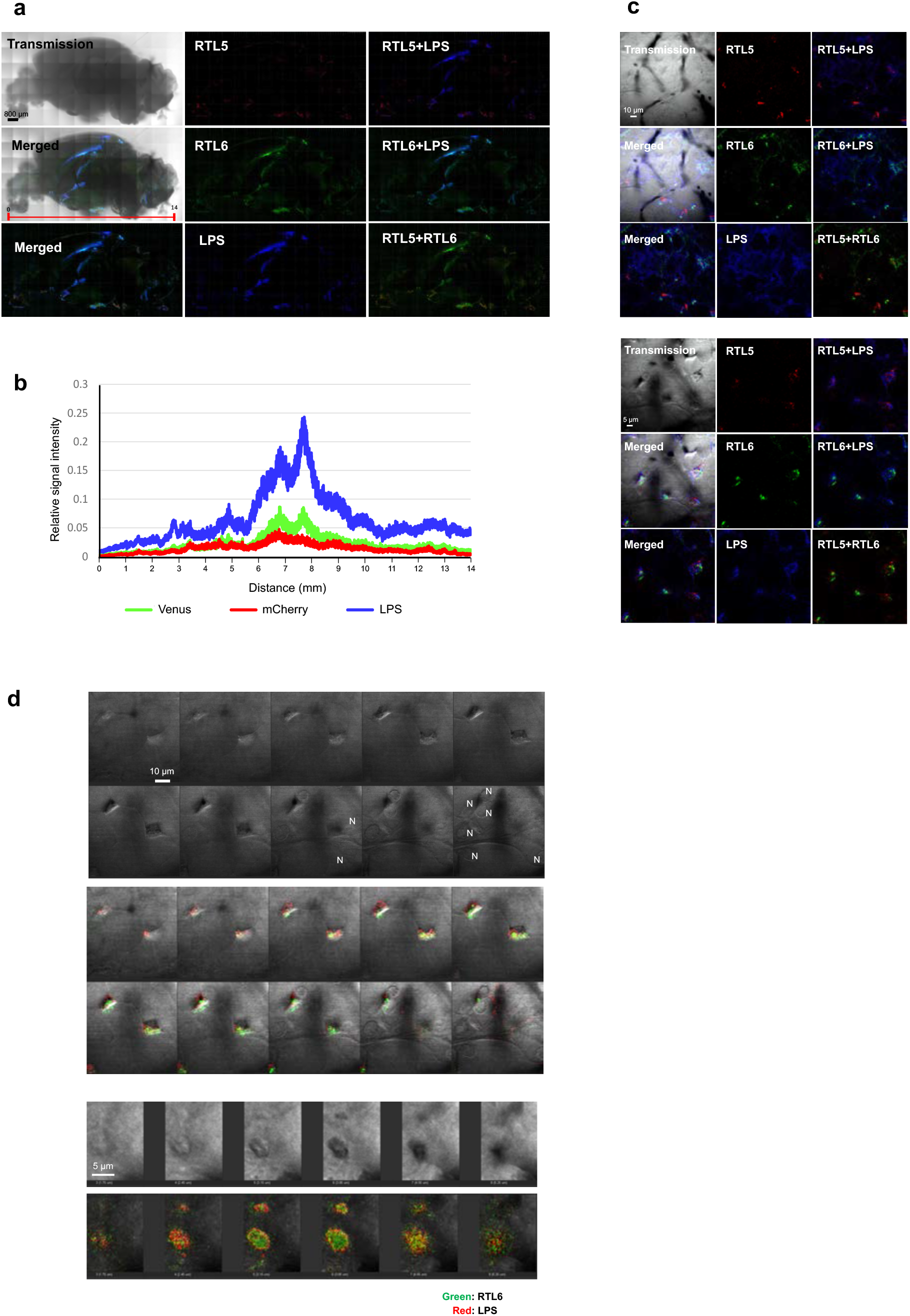
The differential responses of RTL5 and RTL6 to LPS. **(a)** The case of LPS (N=4). RTL5 (red), RTL6 (green) and LPS (blue) signals on the inner face of the brain hemispheres (5w). **(b)** Relative intensity of each RTL5-mCherry (red), RTL6-Venus (green) and LPS (blue) signal/transmission signal is presented along the x-axis (direction from the olfactory bulb to the cerebellum) as shown by a red line in the left middle panel in Fig. 5a. **(c)** Top panels: giant flattened cells along with the blood capillaries in the cerebral cortex. Bottom panels: enlarged pictures showing the large RTL5/RTL6/LPS complexes. **(d)** A large RTL6/LPS complex observed in the giant flattened microglial cells. Two sets of sequence of photographs at 0.7 μm intervals are presented. Top and middle panels: localization of the large RTL6/LPS complex at an intersection of giant flattened cells. Transmission (upper) and fluorescent images (lower). N: nucleus. Green: Venus. Red: Alex594-labelled LPS. Bottom panels: Enlarged transmission (upper) and fluoresecent images (lower) of the large RTL6/LPS complex.

Both RTL6 and RTL5 accumulated at the poly (I:C), a synthetic dsRNA analog, injection sites (shown in artificial blue) in proportion to the amount of Rhodamine-labeled-Poly(I:C) (emission peak 576 nm) (Figs. 7a and b). On higher magnification, the major portion of the injected dsRNA analog had accumulated in the proximity of the nucleus of the microglia and formed a large, chain-like (10 μm long) complex (cyan) with the RTL6 protein (Fig. 7c). RTL5 (red) was also included in the complex (Fig. 7c).

**Figure 7.**
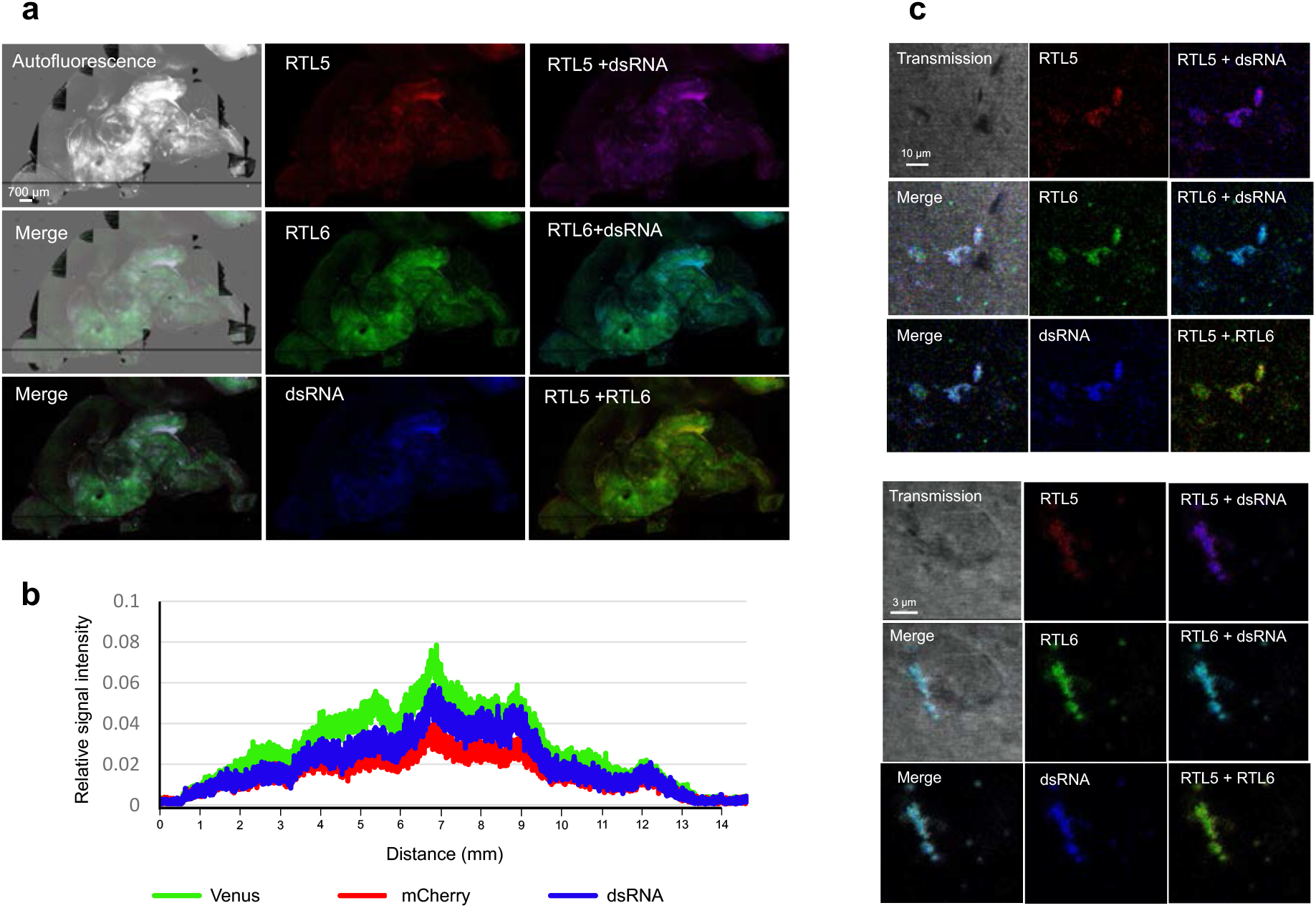
The differential responses of RTL5 and RTL6 to dsRNA. **(a)** The case of dsRNA analog (N=2). RTL5 (red), RTL6 (green) and dsRNA analog (blue) signals on the inner face of the brain hemispheres. **(b)** Relative intensity of each RTL5-mCherry (red), RTL6-Venus (green) and dsRNA analog (blue) signal in P7 neonate. **(c)** Top and bottom panels: microglial cells with RTL5/RTL6/dsRNA complexes.

In contrast, upon injection of non-methylated dsDNA labeled with Cyanine (Cy)3 (emission peak 570 nm, shown in artificial blue), RTL5 mainly reacted to form the RTL5/dsDNA complex (violet) without RTL6 (Figs. 8a and b). The relative signal intensity of RTL5 (red) exceeded that of the RTL6 case in which a higher dsDNA signal had been observed. The majority of the dsDNA was incorporated within two hours after injection into the cytoplasm of the microglial cells, mostly with bilateral processes, and formed an RTL5/DNA complex without any RTL6 protein (Fig. 8c). It should be noted that the relative signal intensity of RTL6 usually exceeded that of RTL5 throughout the brain (Figs. 8d and e).

**Figure 8.**
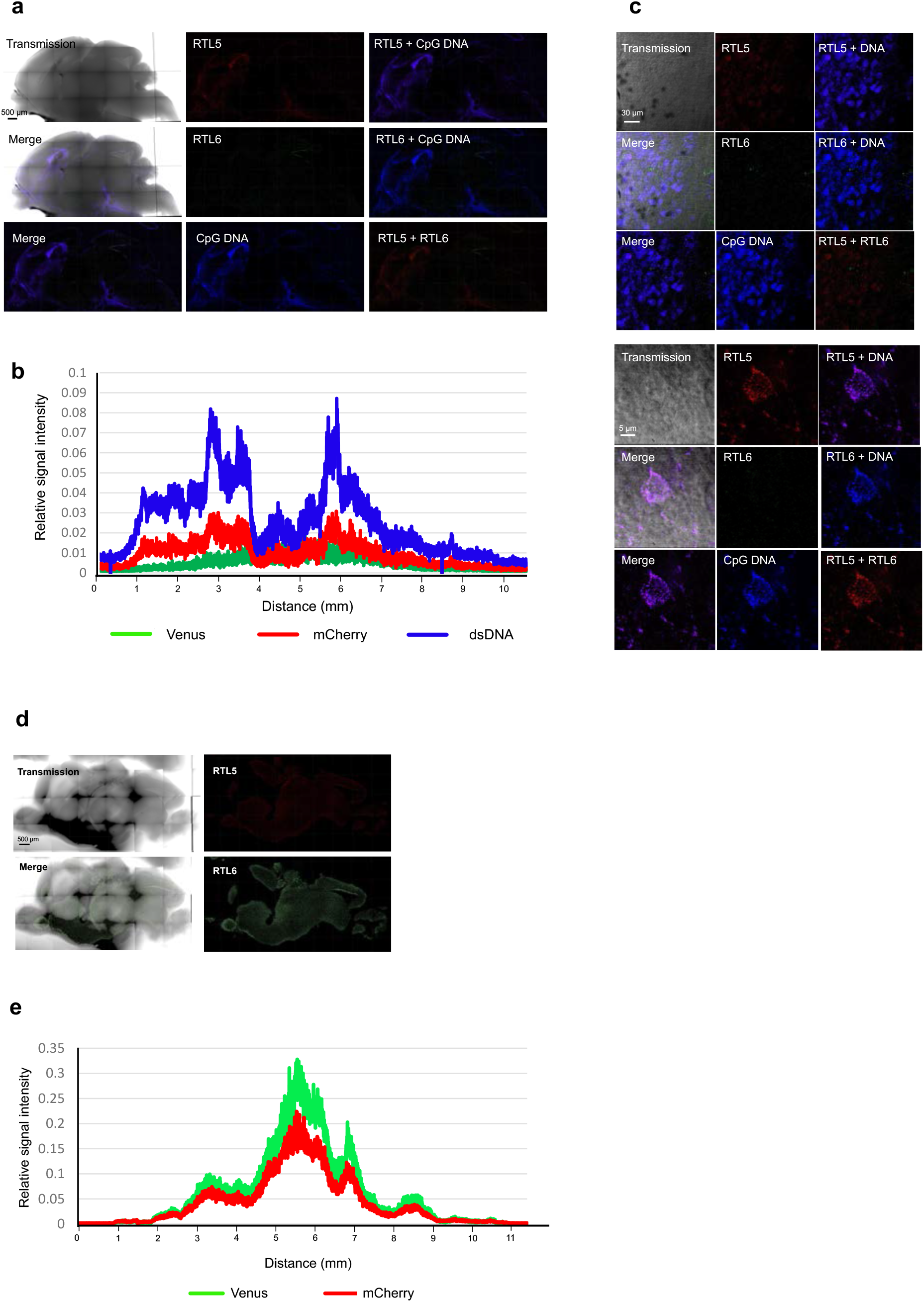
The differential responses of RTL5 and RTL6 to dsDNA. **(a)** The case of non-methylated dsDNA (N=2). RTL5 (red), RTL6 (green) and dsDNA (blue) signals on the inner face of the brain hemispheres. **(b)** Relative intensity of each RTL5-mCherry (red), RTL6-Venus (green) and dsDNA (blue) signal in P3 neonate. **(c)** Top panels: microglial cells with accumulated dsDNA. Bottom panels: microglial cells with RTL5/dsDNA complexes. **(d)** RTL5 and RTL6 expression in non-injection control mouse (*Rtl5-mCherry, Rtl6-Venus* double KI (DKI) mouse (P3 neonate)) (N=3). Top and bottom panels: Transmission, fluorescent and merged images of RTL5 (red) and RTL6 (green), respectively. **(e)** Relative signal intensity of RTL5 and RTL6. In every part of the brain, the relative signal intensity of RTL6-Venus (green) was greater than that of RTL5-mCherry (red) in the DKI mice. Due to the methodological limitation in obtaining the absolute value of each signal intensity, it is not possible to compare the signal intensity between different samples, however, possible to compare the relative intensity of each signal in the same brain samples.

## Discussion

Our data clearly demonstrate that RTL6 and RTL5 function as microglial genes in the front line of innate immunity for the quick clearance of certain pathogens, providing the first evidence that the eutherian-specific genes acquired from retroviral infection function as members of the innate immunity system in the eutherian brain (Figs. 6-8). In other word, eutherian-specific *RTL5* and *RTL6* are also good examples of exaptation (Gould and Vrba, 1982; Brosius and Gould. 1992) because they work as “self constituents” in the Self/Nonself discrimination system. It seems quite likely that, based on the retroviral GAG protein, the leucine zipper motif emerged in the N-terminus and extremely acidic/basic and basic regions of the C-terminus of RTL5 and RTL6 by multiple mutation events (Figs. 1e and f). According to the SWISS-MODEL prediction (Bienert et al., 2017) the last half (108-215 aa) of RTL6 has 6 short helix structures (sH1-6), as does the original GAG protein (Fig. 9).

**Figure 9.**
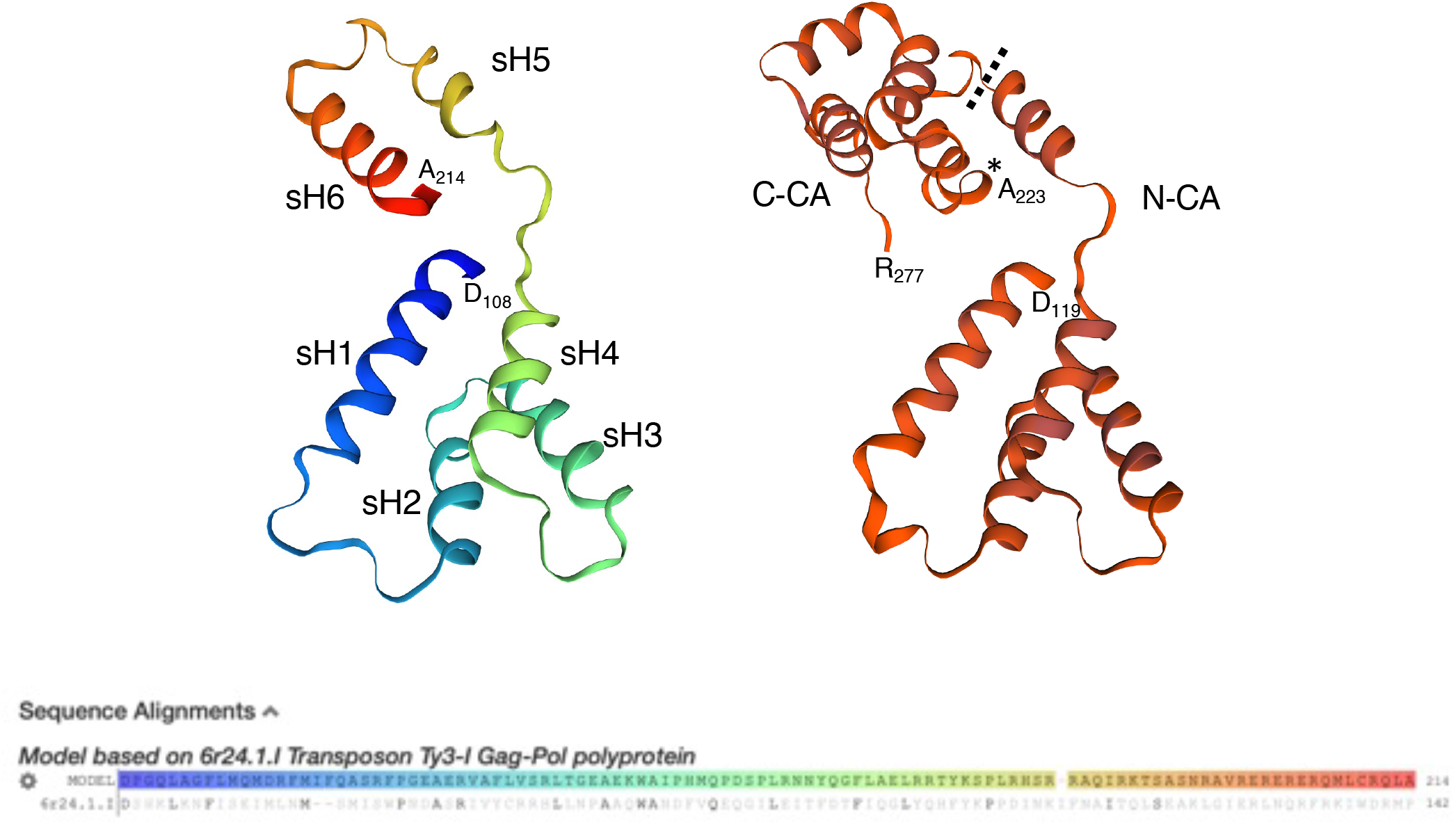
Estimated 3D structure of RTL6 protein. From the entire RTL6 aa sequence (1-243 aa residues), the SWISS-MODEL^19^ predicts that the C-terminus (108-214 aa residues) forms six short helix (sH) structures with strong basic properties in sH5 and sH6. The estimated 3D structures of RTL6 (left) and sushi-ichi GAG proteins (right) are compared. Both of the 3D structures are based on the 6r24.1.1. Transposon Ty3-I Gag-Pol polyprotein.

Interestingly, sH5 and sH6 are strongly basic (pI values: 12.30 and 10.14, respectively), so may play an essential role in the trapping of acidic substances such as LPS and dsRNAs. The dN/dS ratio of *RTL6* (< 0.05) lies between the average dN/dS ratio of the house-keeping genes (approximately 0.093) and that of the Histone H3 gene (< 0.01), one of the most widely conserved genes (Kimura, 1986; Zhang and Li, 2004). This indicates that *RTL6* has been powerfully conserved in eutherians, suggesting that RTL6’s role in LPS removal is vitally important, because LPS is an extremely dangerous pathogen.

Microglia express a variety of TLR proteins, i.e. TLR3, 4 and 9 for dsRNA, LPS and non-methylated CpG DNA, respectively (Fiebich et al., 2018). At the moment, it remains unknown how RTL5 and RTL6 are related to the TLR system in innate immunity. As both proteins exist as intra-as well as extracellular granules, it is possible that they work as guardians standing by for clearance of invaded pathogens independently from the TLR systems. Alternatively, they may also work as sensors of these pathogens in the TLR systems. Recently, there reported two TLR4-independent innate immune responses, such as transient receptor potential (TRP) channels and caspase 4/5 against intracellular LPS (Kayagaki et al., 2013; Meseguer et al., 2014; Shi et al., 2014; Masgaeen and Gurung, 2020). Therefore, redundant pathways may exist in mammals, especially against LPS. Detailed genetic and biochemical analyses of these genes and proteins will be of special interest in unraveling the uniqueness of the present day eutherian innate immune system.

In the neonatal brain microglia are involved in shaping neuronal circuits during development via regulating neurogenesis. They induce filopodia formation by direct contact with neurons and phagocytose supernumerary or unwanted synapses as well as pruning excess astrocytes in the developing amygdala (Hanisch and Kettenmann, 2007; Sierra et al., 2010; Reemst et al., 2016). RTL5 and RTL6 thus seem likely to play an important role in maintaining a clean environment in the developing brain by removing hazardous substances leaking from damaged neuronal cells during neural network formation.

It is known that neuronal cells secrete the activity-regulated cytoskeletal (ARC) protein, a GAG-derived capsid-like substance, for communicating between and among neuronal cells via binding and delivering mRNA (Ashley et al., 2018; Pastuzyn et al., 2018). Recently, the PEG10 and RTL1 proteins were also shown to be able to form a virus-like structure with the ability to deliver mRNA, much like exosomes (Segel et al., 2021). It is deeply interesting to consider that the basic structure of the GAG protein has been in a wide range of applications during evolution. *ARC* was domesticated independently in tetrapods and insects (Ashley et al., 2018; Pastuzyn et al., 2018). *PEG10* in therians and *RTL1, RTL5* and *RTL6* in eutherians, respectively. Therefore, it is quite probable that there are additional brain genes from other retroviral GAG sequences not only in eutherians but other organisms, because the domestication of such genes has proven to be of great benefit.

Among the 11 *RTLs, RTL5* and *RTL6* are the first examples of genes functioning in yolk sac-derived microglia (Figs. 3a, 4e and 5e). Microglia originate from the extraembryonic yolk sac in early development, migrate to the embryo and settle in the brain in the fetal stage, then ultimately propagate throughout the brain over the course of life (Ginhoux et al., 2010, 2013). We previously demonstrated that *PEG10, RTL1* and *LDOC1*, play different but essential roles in the placenta (Ono et al., 2006, Kagami et al., 2008; Sekita et al., 2008; Naruse et al. 2014), another extraembryonic tissue, using knockout mice. Endogenous retroviruses (ERVs) and retrotransposons are usually completely repressed in the fetus while nevertheless being constantly transcribed in the extraembryonic tissues at a low level due to the lower DNA methylation level in these tissues (Kaneko-Ishino and Ishino, 2012, 2015). Therefore, ERV-derived genes could be functionally selected under certain circumstances over the course of evolution extraembryonic tissues at a low level due to the lower DNA methylation level in these tissues (Kaneko-Ishino and Ishino, 2015). In other words, the extraembryonic tissues might have served as a cradle or incubator for retrovirus-derived genes. This may be consistent with and/or complement the recently reported finding of the placenta serving as a dumping ground for genetic defects (Coorens et al., 2021), because the placenta is thus able to tolerate major genetic and/or developmental flaws which affords a tremendous advantage for the survival of the fetus. Our work indicates a novel role of yolk sac in functional evolution of the innate immune system in eutherians.

## MATERIALS AND METHODS

### Mice

All the animal experiments were reviewed and approved by Institutional Animal Care and Use Committee of RIKEN Kobe Branch and Tsukuba Branch, Tokai University and Tokyo Medical and Dental University (TMDU) and were performed in accordance with the RIKEN Guiding Principles for the Care and Use of Laboratory Animals, and the Guideline for the Care and Use of Laboratory Animals of Tokai University and TMDU.

### Comparative genome analysis

The sushi-ichi GAG (AAC33525.1) and mouse RTL5 (NP_001265463.1) and RTL6 (NP_808298.2) protein sequences were obtained from NCBI and Ensemble. Amino acid identity and similarity were calculated using the EMBOSS Water program (http://www.ebi.ac.uk/Tools/psa/emboss_water/) in the default mode. The orthologues of *RTL5* and *RTL6* were identified by a search of the NCBI Gene database (http://www.ncbi.nlm.nih.gov/gene/) using *RTL5* (and *RGAG4*) and *RTL6* (and *LDOC1L*) as the query term. Genomic homology analysis was performed using the mVISTA LAGAN program (http://genome.lbl.gov/vista/mvista/submit.shtml).

The sequences used for *RTL5* comparative genome analysis in Fig. 1a were as follows: Chicken (*Gallus gallus*): NC_052535.1[c2173485-1600210]; Platypus (*Ornithorhynchus anatinus*): NC_041733.1[17047115-18009974]; Echidna (*Tachyglossus aculeatus*): NC_052071.1[c42206678-41135833]; Opossum (*Monodelphis domestica*): NC_008809.1[c72655043-72430642]; Mouse (*Mus musculus*): NC_000086.8[100676255-101558086]; Human (*Homo sapiens*): NC_000023.11[71526681-72577989]; Chimpanzee (*Pan troglodytes*): NC_036902.1[66945448-68002204]; Cattle (*Bos taurus*): NC_037357.1[c79138511-78100425]; Dog (*Canis lupus familiaris*): NC_051843.1[56788745-57634597]; Cat (*Felis catus*): NC_018741.3[60765912-61585357] ; Horse (*Equus caballus*): NC_009175.3[56412010-57205718]; African savanna elephant (*Loxodonta Africana*) NW_003573444.1[c31006766-29638658].

We obtained the *NHSL2* genomic sequences in Fig 1b from the NCBI database and Ensemble. The sequences used for analysis were as follows: Chicken (*Gallus gallus*): bGalGal1.mat.broiler.GRCg7b, XP_015133857.1; Platypus (*Ornithorhynchus anatinus*): mOrnAna1.pri.v4, XP_028923216.1; Opossum (*Monodelphis domestica*): MonDom5, XP_007507794.1; Mouse (*Mus musculus*): GRCm39, XP_011245801.1; Human (Homo sapiens): GRCh38.p13, NP_001013649.2; Chimpanzee (*Pan troglodytes*): Clint_PTRv2, XP_016798720.2; Dog (*Canis lupus familiaris*): ROS_Cfam_1.0, XP_038306208.1; Cat (Felis catus): Felis_catus_9.0, XP_019679523.1; African savanna elephant (*Loxodonta Africana*): Loxafr3.0, XP_023407160.1; Armadillo (*Dasypus novemcinctus*) Dasnov3.0, XP_023439655.1+XP_023439656.1; Sloth (*Choloepus didactylus*): mChoDid1.pri, XP_037677403.1.

For *RTL6*, we obtained the *SHISAL1*-*PRR5* genomic sequences in Fig. 1c from the NCBI database. The sequences used for analysis were as follows: Chicken (*Gallus gallus*): NC_006088.4[69115783-70071731]; Platypus (*Ornithorhynchus anatinus*): NC_041741.1[c52020074-51756772]; Echidna (*Tachyglossus aculeatus*): NC_052079.1[c58674342-58403296]; Opossum (*Monodelphis domestica*): NC_008808.1[15497440-16650585] and NW_001583545.1[1-415929]; Tasmanian devil (*Sarcophilus harrisii*): NW_003843556.1[1-720570] and NW_003844018.1[1-136229]; Mouse (*Mus musculus*): NC_000081.6[84153625-84874824]; Human (*Homo sapiens*): NC_000022.11[43913843-45028826]; Dog (*Canis lupus familiaris*): NC_006592.3[21015698-21743182]; African savanna elephant (*Loxodonta Africana*) NW_003573493.1[76766-1008174]; Armadillo (*Dasypus novemcinctus*) NW_004489016.1[1-309452]; Sloth (*Choloepus hoffmanni*) KN190031.1[1-672150].

### Estimation of the pairwise dN/dS ratio

The nonsynonymous/synonymous substitution rate ratio (dN/dS) was estimated by using CodeML (runmode: −2) in PAML (Xu and Yang, 2013). An amino acid sequence phylogenic tree was constructed with MEGA7 (Kumar et al., 2016) using the Maximum Likelihood method based on the JTT matrix based model. The codon alignment of cDNA was created with the PAL2NAL program (www.bork.embl.de/pal2nal/) (Suyama et al., 2006).

The *RTL5* genome sequences used for the dN/dS analysis (Table 1) were the following: Mouse: NC_000086.8[c101114468-101112669]; Human: NC_000023.11[c72131540-72129831]; Chimpanzee: NC_036902.1[c67554437-67552728]; Dog: NC_051843.1[c57238370-57236637]; Cattle: NC_037357.1[78545112-78546836]; Horse: NC_009175.3[c56844294-56842597]; Elephant: NW_003573444.1[30233756-30235345]; Manatee: NW_004444006.1[c9027102-9025528]; Armadillo: NW_004489802.1[c991100-989476]; Sloth: KN181417.1[c16728-15101]. The *RTL6* genome sequences used for analysis were the following: Mouse: NC_000081.6[84556462-84557193]; Human: NC_000022.11[44496837-44497556]; Chimpanzee: NC_006489.4[31270844-31271566]; Dog: NC_006592.3[21316309-21317028]; Cattle: AC_000162.1[115850420-115851139]; Horse: NC_009171.2[40642984-40643703]; Elephant: NW_003573493.1[337200-337919]; Manatee: NW_004444005.1[6540826-6541545]; Armadillo: NW_004489016.1[146135-146854]; Sloth: KN190031.1[86985-87704].

### Rapid Amplification of cDNA Ends (RACE)

For the 5’-RACE experiment, the 5’-Full RACE Core Set (TaKaRa) was used to extend *Rtl6* mRNA from the mouse brain at 8 weeks of age, according to the manufacturer’s instructions. The first strand cDNA synthesis was carried out with 1 μg of total RNA using Rtl6 5’-RACE GSP (Fig. 2a): 5’-AGGGTGTCAACTACG-3’ (5’-phospholylated). The 5’-RACE PCR was performed with Ex*Taq* polymerase (TaKaRa) using the following primers: Sirh3-race-F1: 5’-CTGAAAGCCCAGCCTCTGCC-3’ and Rtl6-race-R1: 5’-TGGAGGTCCGAGGTTGGACC-3’. The 5’-RACE PCR products (1st 5’-RACE products) were confirmed by 1.5 % agarose gel electrophoresis. Nested PCR was performed with Ex*Taq* polymerase using the following primers: Sirh3-race-F2: 5’-TGGTGCCAGCGCTCAGATGG-3’ and Rtl6-race-R2: 5’-AGGTTGGACCATGCTGGCGG-3’. A 1/50 dilution of the 1st 5’-RACE products was used as a template. The nested PCR products were separated by 1.5 % agarose gel electrophoresis and extracted from the gel. The extracted PCR products were cloned into a pGEM-Teasy vector (Promega). The DNA sequence was determined by Sanger sequencing. For the 3’-RACE experiment, the 3’-Full RACE Core Set (TaKaRa) was used to extend *Rtl6* mRNA from the mouse brain at 8 weeks of age according to the manufacturer’s instructions. First strand cDNA synthesis was carried out with 1 μg of total RNA. The 3’-RACE PCR was performed with Ex*Taq* polymerase (TaKaRa) using the following primers: Rtl6 3’-RACE GSP: 5’-ATCCAGCCTCCAACGGGACC-3’ and 3 sites Adaptor primer: 5’-CTGATCTAGAGGTACCGGATCC-3’. The 3’-RACE PCR products (1st 3’-RACE products) were confirmed by 1 % agarose gel electrophoresis. Semi-nested PCR was performed with Ex*Taq* polymerase using the following primers: Rtl6-race-F4: 5’-TCCAACGGGACCAATCCCGC-3’ and 3 sites of adaptor primer, with a 1/50 dilution of the 1st 3’-RACE products as a template. The semi-nested PCR products were separated by 1 % agarose gel electrophoresis and extracted from the gel. The extracted PCR products were cloned into a pGEM-Teasy vector (Promega). The DNA sequence was determined by Sanger sequencing.

### Generation of the *Rtl6-Venus* knock-in mice

The *Rtl6-Venus* fusion protein construct (pRtl6CV) was generated by Gateway cloning technology (Thermo Fisher Scientific) and the Kuroyanagi *et al*. method (Kuroyanagi et al., 2010). The PCR fragment, including the Rtl6 5’-UTR to ORF (the Rtl6N fragment), was generated using PrimeSTAR Max DNA Polymerase (TaKaRa) and the following primers: Rtl6attB1: 5’-GGGGACAAGTTTGTACAAAAAAGCAGGCTCAACCGAAGGATGAGAGGGTC-3’ and Rtl6NattB5r: 5’-GGGGACAACTTTTGTATACAAAGTTGTCCGAGGTTGGACCATGCTGGCG-3’. The PCR fragment, including the Rtl6 5’-UTR to Sirh3 ORF end (the Rtl6C fragment), was generated using PrimeSTAR Max DNA Polymerase (TaKaRa) and the following primers: Rtl6attB1 and Rtl6CattB5r: 5’-GGGGACAACTTTTGTATACAAAGTTGTAAGGTTCCGGCCACGAGAGGGCA-3’.

The pDONRRtl6N and pDONRRtl6C vectors were constructed by the Gateway BP reaction using the following fragments and vectors: pDONRRtl6N: the Rtl6N fragment and pDONR221 P1-P5r (Thermofisher), pDONRRtl6C: the Rtl6C fragment and pDONR221 P1-P5r. The Rtl6-3’UTR fragment was obtained by PCR amplification using the following primers: Rtl6XhoI-F 5’-CGCCTCGAGGGACTTGCCACCACCCTGGTAG-3’ and Rtl6EcoRI-R 5’-CGCGAATTCCTCCTGTCCTGGTCTTGCAAAGG-3’. The Rtl6-3’UTR fragment was ligated to the *Xho*I and *Eco*RI digested pBluescript SK(+) vector. The inverse-PCR fragment of Rtl6-3’UTR was amplified using the following primers: Rtl6attB1: 5’-GGGGAGCCTGCTTTTTTGTACAAACTTGTCCGGTACCCAATTCGCCCTATAG-3’ and Rtl6attB2: 5’-GGGGACCCAGCTTTCTTGTACAAAGTGGTCGGACTTGCCACCACCCTGGTAG-3’. The pDONRRtl6-3’UTR vector was constructed by the Gateway BP reaction using pDONR221 P1-P2 (Thermofisher) and the Rtl6-3’ UTR fragment. The pRtl6NV and pRtl6CV vectors were constructed by the Gateway LP reaction using the following vectors: pRtl6NV: pDONRRtl6N, pDONRRtl6-3’UTR and pENTR-L5-Venus-L2, pRtl6CV: pDONRRtl6C, pDONRRtl6-3’UTR and pENTR-L5-Venus-L2.

To generate the *Rtl6* KI targeting vector, we obtained two PCR fragments, the 5’-arm (6.5 kb) and 3’-arm (3.5 kb), using PrimeSTAR Max DNA Polymerase (TaKaRa). The C57BL/6N genome was used as the PCR template. For Rtl6 5’- and 3’-arm cloning, we used the following primers: 5’-arm: Rtl6KI-LA-F1: 5’-CTGACTgtcgaccaattgCCTGCTGTTTGGTGGTTGAGCCTCTG-3’, Rtl6KI-LA-R1: 5’-CTGACTggattcCTTTACGATTCCTACCCAGGCCGCTC-3’, Rtl6KI-LA-F2: 5’-CTGACTgtcgacGGGAATGTAGAGGCAGGAGAGGTTCAAGG-3’ and Rtl6KI-LA-R2: 5’-CTGACTgcggccgcttaattaaGGAGTGTTCCAGGAGCTGAGTATCCGTG-3’; 3’-arm: Rtl6KI-SA-F1: 5’-CTGACTgtcgacCTACAGCTCTTGCTGCCCCAGGC-3’ and Rtl6KI-SA-R1: 5’-CTGACTgcggccgcGTGTGGGCTGAAGACAGGTGGGTTG-3’. The middle arm (1kb: Rtl6NV, 1.5 kb: Rtl6CV) fragments were generated by restriction enzyme digestion of pRtl6NV and pRtl6CV, respectively. All of the arm fragments were inserted into a pNT1.1 vector.

The establishment of knock-in ES cells and generation of chimeric mice are conducted as previously described (Fujihara et al., 2013). In brief, EGR-G101 ES cells were electroporated with linearized DNA, and then screened by PCR after positive/negative selection. Chimeric mice were produced by the 8-cell microinjection method. To remove the flox region, we injected a pCAG/NCre plasmid (Sato et al., 2000) into the fertilized eggs generated by in vitro fertilization (IVF) from C57BL/6N eggs and Rtl6-CV mutant sperm.

### Immunoprecipitation and Western blotting

Adult brain (8 w) was dissected into seven parts and each part was powderized in liquid N_2_ using a Multi-beads shocker (MB1050, YASUI KIKAI, Osaka). The powder samples of WT and RTL6-CV cerebrum (46 mg and 48.1 mg, respectively) were dissolved in 150 μl of RIPA buffer, 50 mM Tris-HCl (pH 8.0), 150 mM NaCl, 0.5 % Sodium deoxycholate, 0.1 % SDS, 1 % NP-40 (IGEPAL CA-630) and 1 mM EDTA, supplemented with 20× protease inhibitor solution (Sigma-Aldrich, P2714) on ice for 30 min. After 20 min of centrifugation (10,000g, at 4°C), the supernatant was mixed with anti-GFP (RatIgG2a), Monoclonal (GF090R), CC, Agarose Conjugate (NACALAI TESQUE) and incubated overnight at 4°C. The agarose beads were washed four times with 500 μl of RinseBuffer, 50 mM Tris-HCl (pH 8.0), 150 mM NaCl at 4°C. Then the beads were incubated with 60 μl of SDS sample buffer and directly applied to gel electrophoresis using a 10% acrylamide gel. Western blot analysis was performed using a standard protocol. After blotting on a Hybond-P (GE Healthcare) membrane, the Sirh3-Venus fusion protein was detected with an ECL Prime Western Blotting Detection kit (GE Healthcare) using an anti-GFP antibody (MBL, Code No. 598) and an anti-Rabbit Goat Immuno-globulins/HRP (DAKO, P0160) as 1^st^ and 2^nd^ antibodies. Signals were detected on an AE-9300 Ez CaptureMG (ATTO, Tokyo).

### Quantitative RT-PCR

Total RNA was prepared from frozen tissues using ISOGEN (NIPPON GENE) and ISOGEN-LS (NIPPON GENE). The cDNA was made from total RNA (1 μg) using Revertra Ace qPCR RT Master Mix (TOYOBO). Quantitative RT-PCR analysis was performed using Fast SYBR Green Master Mix (Life technologies) and a StepOnePlus System (ABI) by means of an absolute quantification method. Student’s *t*-test was used for statistical analysis. The following primer sequences were used: Gapdh-F: 5’-CACTCTTCCACCTTCGATGC-3’ and Gapdh-R: 5’-CTCTTGCTCAGTGTCCTTGC-3’; Rtl6-F2: 5’-GTGTTGGGTGGCAAATGCTCGG-3’ and Rtl6-R2: 5’-GGACCTCCCAGACACTGCAAGC-3’.

### Imaging using Confocal Laser Scanning Fluorescence Microscope

Fresh brain and brain slices (2 mm in depth) from *Rtl5*-CmCherry and *Rtl6*-CVenus KI mice were used for analysis with a ZEISS LSM880 (ZEISS, Germany) with and without fixation using 4 % paraformaldehyde (PFA). Samples were covered with 10% glycerol solution for protection from drying. The samples were observed using a Plan-Apochromat lens (10x, numerical aperture =0.45, M27, ZEISS). The tiling with lambda-mode images was obtained using the following settings: pixel dwell: 1.54 μs; average: line 4; master gain: ChS: 1250, ChD: 542; pinhole size: 33 μm; filter 500 – 696 nm; beam splitter: MBS 458/514; lasers: 514 nm (Argon 488), 0.90 %. For the tiling-scan observations, the tiling images were captured as tiles: 84, overlap in percentage: 10.0, tiling mode: rectangular grid, in size: x; 15442.39 μm, y; 9065.95 μm. Spectral unmixing and processing of the obtained images were conducted using ZEN imaging software (Carl Zeiss Microscopy, Jena Germany). The spectrum from Venus proteins (Maximum peak emission fluorescence wavelength: 528 nm) was only detected in the samples from *Rtl6*-CV and *Rtl5-CmC* and *Rtl6*-CV double KI mice, not the wild type control samples. The mCherry signal (Maximum peak emission fluorescence wavelength: 610 nm) detected in *Rtl5-CmC* and *Rtl5-CmC* and *Rtl6*-CV double KI mice was distinguished from Af610 nm using the peak shape, such as the width and/or co-existence of a second peak. The relative intensity of RTL5-mCherry (red), RTL6-Venus (green) and LPS (blue) signals along the x-axis of the brain (from the olfactory bulb to cerebellum regions) was calculated from 3D scanning data. The total intensity of each signal on and above each y-axis was summed, divided by the transmission signal and presented as the relative signal intensity on the y-axis in this figure.

### Immunostaining

For immunostaining, fresh frozen optimal cutting temperature (OCT) compound (Tissue-Tek, Sakura, FineTek)-embedded brain sections were fixed in 4% PFA (26126-25, Nacalai Tesque) in 0.1 M PBS at RT for 20 min and washed 3 times with PBS for 5 min. Then they were treated with PBS containing 0.2% Triton-X100 for 20 min and washed 3 times with PBS for 5 min. After the blocking reaction using PBS containing 5 % normal goat serum and 1% BSA at RT for 30 min, the sections were first incubated with an anti-Iba1 rabbit antibody (MBL, Code No. D513-A48). After being washed 3 times with PBS for 5min, incubated with an anti-rabbit-Alexa Fluor 488 antibody (MBL, Code No. D533-A62) at 4°C overnight and finally washed 3 times with PBS for 5min, the images were captured with a ZEISS LSM880 (ZEISS, Germany).

### Generation of the *Rtl5-mCherry* knock-in mice

*Rtl5-CmCherry* mouse was generated by pronuclear microinjection of CRISPR/Cas system (see Fig. 5a), essentially as in a previous report (Aida et al., 2015), using single-stranded (ss) donor DNA. ssDNA was prepared by 5’phosphorylated primer-mediated PCR from the template plasmid followed by digestion of phosphorylated strand with lambda exonuclease. The template plasmid for the mCherry-targeting ssDNA was constructed with 1.5 kb long 5’ and 3’ arms amplified from the C57BL/6N genome using PrimeSTAR GXL DNA Polymerase (TaKaRa), the upstream genome sequence of the stop codon of Rtl5 and downstream of the predictive cut site by Cas9, respectively. The C-terminus of Rtl5 was fused with mCherry by means of a cloning enzyme (In-Fusion® HD Cloning Kit, Takara Bio) and it was inserted into a pGEM T Easy vector (Promega). A T to A silent mutation was introduced into the position of *Rtl5* Threonine 593 (ACT to ACA) to inhibit recutting of genomic DNA after proper editing by CRISPR/Cas system. Double-stranded DNA (dsDNA) fragments (1142 base pairs) containing 5’ and 3’ homology arms (233 and 201 base pairs, respectively) and an mCherry coding sequence were amplified from the template plasmid by PCR using PrimeSTAR GLX DNA polymerase (TAKARA) along with the sense and antisense primer pair (5’-ATTACCTGGGGGTATCCCCTT-3’ and 5’-CACTCTTCTGGTTGTGGTTGC-3’). The antisense primer was pre-phosphorylated at its 5’ end (Fasmac). Amplified dsDNA fragments were column-purified using a MiniElute PCR Purification Kit (QIAGEN) according to the manufacturer’s recommendations.

Ten μg of the purified dsDNA were treated with Lambda exonuclease (NEB) in 50 μl of reaction solution at 37°C for 30 minutes and then at 75°C for 10 minutes to digest the phosphorylated strand and produce single-stranded DNA (ssDNA). A part of the reaction solution was analyzed for digestion efficiency and ssDNA production by electrophoresis using 1% agarose gel. Total ssDNA was gel-electrophoresed following exonuclease treatment and quality assessment, and gel-purified using a Long ssDNA Preparation Kit for 3 kb (Biodynamics Laboratory Inc.) according to the manufacturer’s recommendations (∼2 μg of ssDNA was produced from 10 μg of amplified dsDNA). Purified ssDNA was stored at 4°C until required, assessed for its quality by Sanger sequencing and injected into mouse pronuclei at final concentration of 5 ng/μl within 1 week of preparation. Just before pronuclear injection, ssDNA was mixed with other components of the CRISPR/Cas system (Cas9 protein, crRNA and tracrRNA).

### LPS, dsRNA, and non-methylated DNA injection to the brain

*Rtl5*-CmC and *Rtl6*-CV double hetero mice (P2 neonates to 5w young adults) were used for the injection experiments after being anesthetized using isoflurane. Ten∼20 μl of Alexa488-labeled LPS (MBL, Code No. D558-A32), Rhodamine-labeled dsRNA (MBL, Code No. D488-A24), and Cyanogen 3-labeled nonmethylated dsDNA (see below) injected using 20 ng/μl solution, respectively, and 1ml insulin syringes and a 26 G needle. One min after the injection, the needle was pulled out and kept out for 10 min, then the fresh brain was dissected out in cold PBS solution. The inner surface of the brain hemispheres was analyzed with a ZEISS LSM880 (ZEISS, Germany) before and after fixation with 4% PFA. Cyanogen 3-labeled nonmethylated dsDNA was made by mixing two complementary oligo DNAs, heating at 60 °C for 5 min and annealing at RT. The oligo DNAs used were 5’-Cyanogen 3 labelled GACGTTGACGTTGACGTTGACGTT and 5’-Cyanogen 3-labeled-AACGTCAACGTCAACGTCAACGTC.

## Author contributions

M.I., J.I., A.M., F.I. and T.K.-I. performed the experiments and analyzed the data. I.M. and M.I. generated *Rtl6*-CV mice and A.M., T.S. and Y.H. generated *Rtl5*-CmC mice. F.I. and T.K.-I. designed the study and F.I., and T.K.-I. wrote the manuscript. All authors agree to be accountable for the content of the work.

## Acknowledgements

We thank N Takayasu and T Umegaki of Tokai University for technical assistance and breeding the mice, H Kuroyanagi of TMUD for his providing a pENTR-L5-Venus-L2 vector, Y Ito of TMDU for his advice on estimation of the RTL6 structure using SWISS-MODEL and Y Niimura of the University of Tokyo for his advice on pairwise dN/dS analyses and phylogenic analyses. We also thank NPO Biotechnology Research and Development and Tokai University Support Center for Medical Research and Education for technical assistance in generating RTL6-Venus KI mice and Takako Usami of TMDU for technical assistance in making the RTL5-mCherry KI mice. Pacific Edit reviewed the manuscript prior to submission.

## Materials & Correspondence

Material requests should be addressed to.

## Data availability

Accession codes for genome data are provided in the Materials and Methods section.

